# Ciliary chemosensitivity is enhanced by cilium geometry and motility

**DOI:** 10.1101/2021.01.13.425992

**Authors:** David Hickey, Andrej Vilfan, Ramin Golestanian

## Abstract

Cilia are hairlike organelles involved in both sensory functions and motility. We discuss the question of whether the location of chemical receptors on cilia provides an advantage in terms of sensitivity. Using a simple advection-diffusion model, we compute the capture rates of diffusive molecules on a cilium. Because of its geometry, a non-motile cilium in a quiescent fluid has a capture rate equivalent to a circular absorbing region with ~ 4× its surface area. When the cilium is exposed to an external shear flow, the equivalent surface area increases to ~ 10×. Alternatively, if the cilium beats in a non-reciprocal way, its capture rate increases with the beating frequency to the power of 1/3. Altogether, our results show that the protruding geometry of a cilium could be one of the reasons why so many receptors are located on cilia. They also point to the advantage of combining motility with chemical reception.

## I. INTRODUCTION

Cilia are small hairlike organelles with a microtubule-based core structure that protrude from the cell surface. They are found on most eukaryotic cells [1] and can be broadly classified into two categories: primary and motile. Primary cilia, of which there is only one on each cell, have primarily sensory functions (as receptors for chemical, mechanical, or other signals) [2–6]. Due to their shape and their role in signalling, they are often referred to as “the cell’s antenna” [7, 8]. Motile cilia, typically appearing in larger numbers [9, 10], move the surrounding fluid by beating in an asymmetric fashion [11, 12], and often with some degree of coordination [13, 14]. They play a key role in a number of processes, including the swimming and feeding of microorganisms [15, 16], mucous clearance in airways [17], fluid transport in brain ventricles [18], and egg transport in Fallopian tubes. However, there are exceptions to this classification. Primary cilia in the vertebrate left-right organiser are motile and drive a lateral fluid flow that triggers, through a mechanism that is not yet fully understood, a distinct signalling cascade determining the body laterality [19]. There is also mounting evidence that motile cilia can have various sensory roles [20], including chemical reception [21]. Adversely, receptors localized on motile cilia, such as ACE2, can also act as entry points for viruses including SARS-CoV-2 [22]. Some chemosensory systems, including include olfactory neurons [23, 24] and marine sperm cells [25], are known to achieve a sensitivity high enough to detect a small number of molecules.

The sensitivity of a chemoreceptor is characterised by its binding affinity for the ligand, as well as its association/dissociation kinetics. If the time-scale of ligand dissociation is longer than the time-scale of the changes in ligand concentration, the sensitivity is determined by the binding rate alone. Because diffusion is fast on very short length scales, only 1% of the surface area of a cell or cilium needs to be covered in high-affinity receptors to obtain near-perfect adsorption [26]. Even if this condition is not satisfied, the membrane itself could non-specifically bind the ligands with near-perfect efficacy, which then reach the receptors in a two-stage process. In either of these cases, as long as there is no advection, the binding rates can be estimated using the theory of diffusion-limited reactions [27]. This binding rate is known as the diffusion limit, and it has already been shown that flagella-driven swimming microorganisms can break the diffusion limit in order to increase their access to nutrients [28].

The increasingly overlapping functions of sensory and motile cilia lead to the natural question about the advantage of placing receptors on a cilium, or in particular on a motile cilium. Because of its small volume, a cilium forms a compartment that facilitates efficient accumulation of second messengers [4, 7]. Placing receptors on a protrusion, away from the epithelial surface, could have other advantages, like avoiding the effect of surface charges or the glycocalycx. It has also been suggested that the location of chemoreceptors on cilia exposes them to fluid that is better mixed [7]. A recent study suggests that the hydrodynamic interaction between motile and sensory cilia can enhance the sensitivity of the latter [29]. However the question of how the geometry and motility of cilia affect their ability to capture and detect ligands has still remained largely unexplored.

In this paper, we investigate the theoretical limits on association rates of ligands on passive and motile cilia. In particular, we address the question of whether the elongated shape of a cilium and its motility can improve its chemosensory effectiveness. By using analytical arguments and numerical simulations, we show that the capture rate of a cilium is significantly higher than that of a receptor located on a flat epithelial surface. Motile cilia can further improve their chemosensitivity. Finally, we show that a cilium within an immotile bundle has a lower capture rate than an isolated cilium, but a higher one when the cilia are sufficiently motile.

## II. RESULTS

In this study, we calculate the second-order rate constant for diffusive particle capture on a cilium. We discuss scenarios where the fluid and the cilium are at rest, where the fluid exhibits a shear flow, where the cilium is actively beating, and where a bundle of hydrodynamically interacting cilia absorbs particles.

We consider a perfectly absorbing cilium protruding from a non-absorbing surface, in a fluid containing some chemical species with a concentration field *c*. Far from the cilium, the unperturbed concentration has a constant value *c*_0_. The rate constant *k* is defined such that

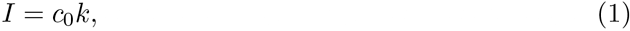

where *I* is the capture rate, defined as the number of captured particles per unit time.

Since the aforementioned cilium is perfectly absorbing, we define an absorbing boundary condition such that the concentration of the chemical species is zero at every point on the cilium’s surface. We assume that the epithelium around the cilium does not absorb particles and it is therefore described with a reflecting boundary condition at *z* = 0. The geometry and boundary conditions are illustrated in Fig. 1.

**FIG. 1:**
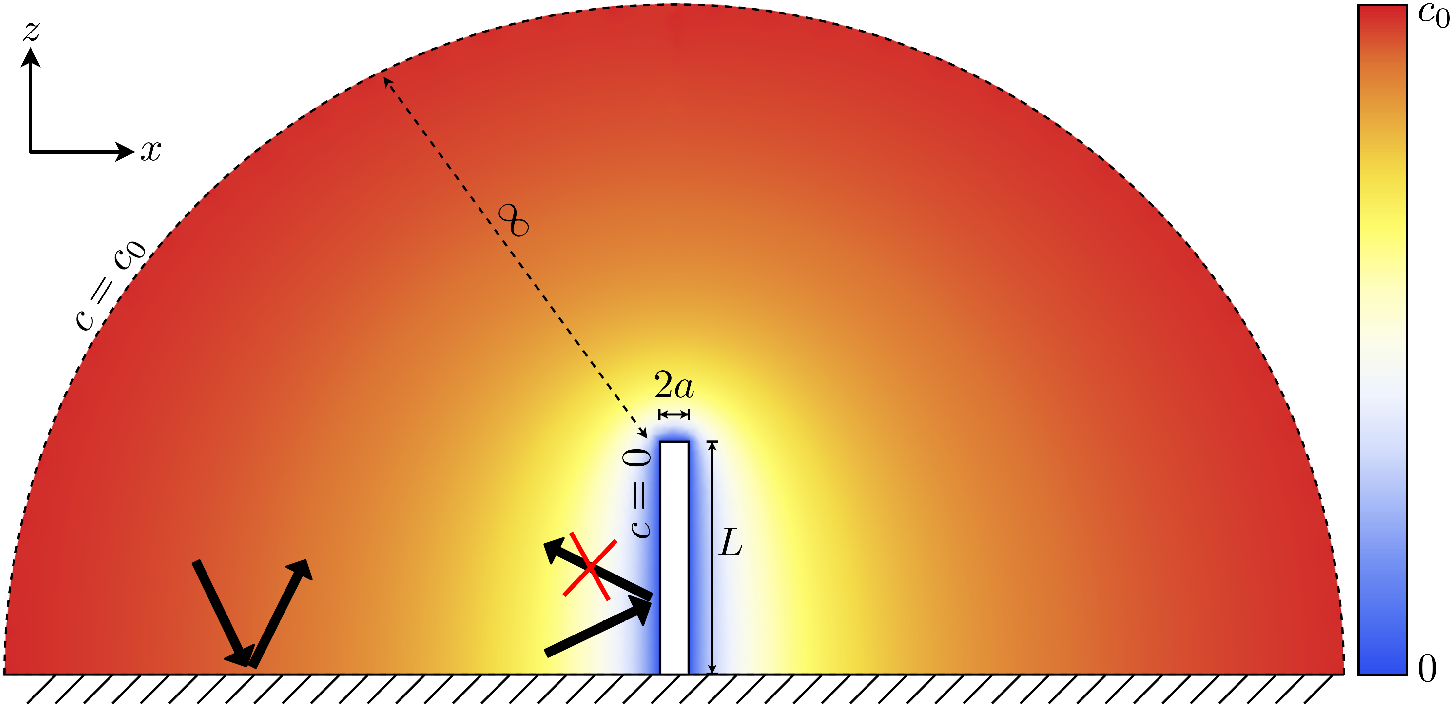
The concentration boundary conditions and general setup of the problem to be solved. The cilium satisfies an absorbing boundary condition, and there is a constant concentration an infinite distance from the cilium. The coloured overlay shows the concentration field in the absence of any fluid flow.

### A. Cilium in quiescent fluid

We consider a cilium (modelled as a cylinder next to a boundary at *z* = 0) in a quiescent fluid, with the goal of determining its capture rate constant in the absence of advection. In the case where there is a steady state with no advection, an analogy to electrostatics can be used to determine the rate constant [26]. The advection-diffusion equation in this case reduces to

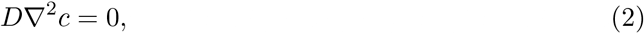

where *D* is the diffusion constant. This is equivalent to the Laplace equation for source-free electrostatics, in which the electrostatic potential *ϕ* obeys

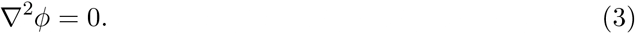

If the boundary conditions in the electrostatic case are chosen such that the body has a potential at its surface of −*V*_0_, the boundary conditions are equivalent as well. The rate constant is determined by the integral of the current density **J** over the surface, which follows from Fick’s law:

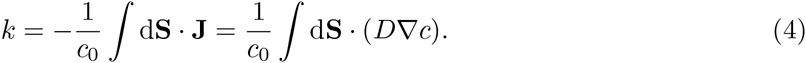

In the electrostatic version of the problem, the equivalent expression for self-capacitance is

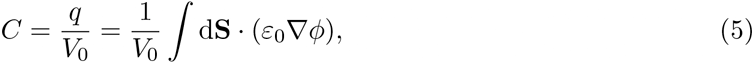

where −*q* is the charge on the body. By analogy, the rate constant can be expressed as:

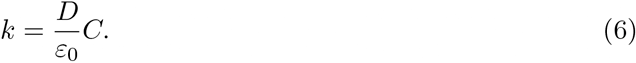

The electrostatic equivalence allows us to translate the calculation of the capture rates to a capacitance problem with a greater number of available solutions in the literature.

There is no closed-form expression for the capacitance of a cylinder, so we loosely approximate this cylinder as half of a prolate spheroid with semi-major axis *L* and semi-minor axis *a*. Using its self-capacitance in the limit *L* ≫ *a* [30], we find the rate constant:

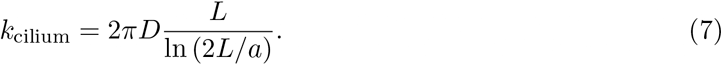

Experiments on olfactory cilia have shown that there is a strong correlation between cilium length and chemosensitivity [31], which is in agreement with this calculation.

To quantify the advantage of localising the receptors on a cilium, we compare it with a case where the receptors form a flat circular patch on the epithelial surface (Fig. 2a). Again, we assume that the receptor patch has a perfectly absorbing surface, while the surface surrounding it is reflective. We determine the size of the patch needed to attain the same rate constant as the cilium. The rate constant for a circular patch on the reflective boundary can be found by applying the electrostatic analogy to the well-known result for the self-capacitance of a thin conducting disc of radius *R*: [26]:

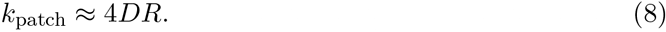

**FIG. 2:**
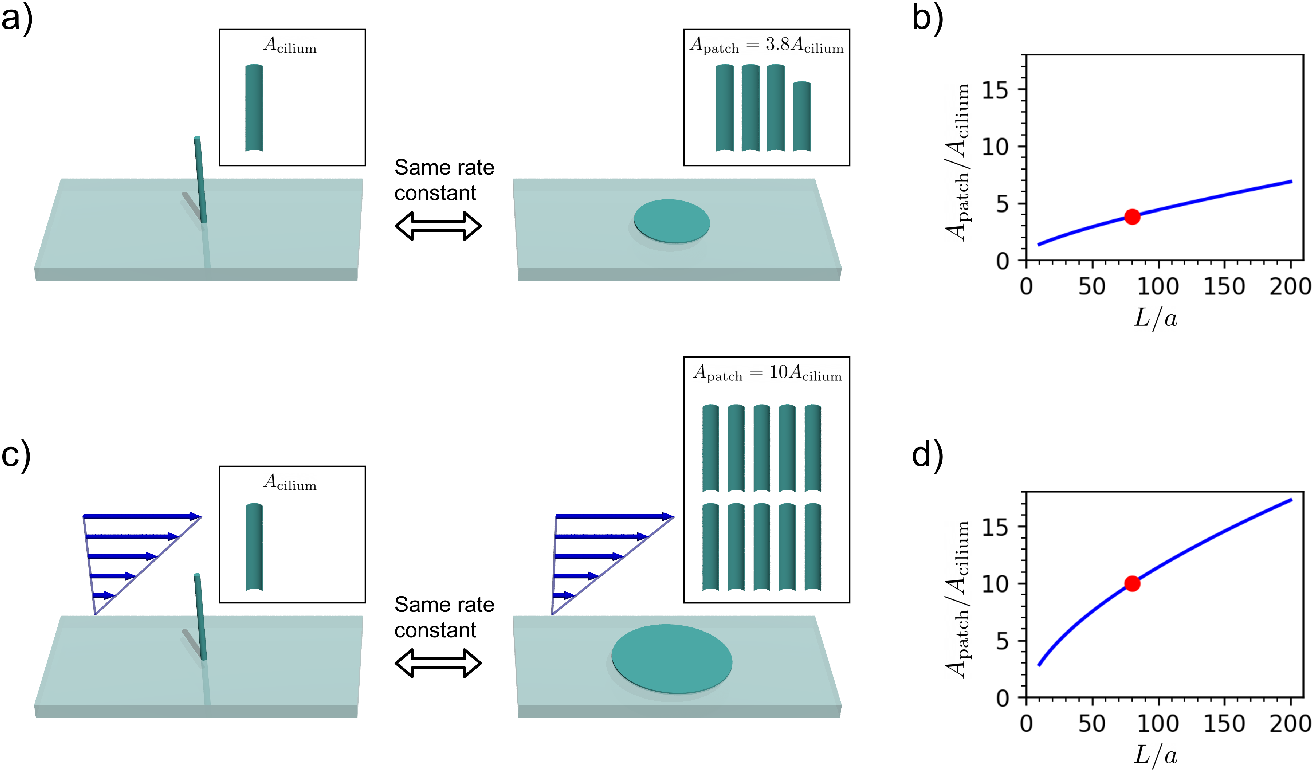
Comparison between capture rates of a non-motile cilium and a circular patch on the surface. All diagrams use *L/a* = 80, indicated on the graphs by a red dot. (a) In a quiescent fluid, the cilium has the same capture rate as a surface patch with 3.8 times the surface area. (b) The area ratio *A*_patch_*/A*_cilium_ as a function of the cilium aspect ratio *L/a* in a quiescent fluid. (c) In a shear flow at a high Péclet number, the capture rate of the cilium reaches that of a surface patch with 10 times the surface area. (d) The area ratio as a function of the aspect ratio in the high Péclet number limit.

We find that the patch has a much larger surface area than the cilium which has the same rate constant. We can calculate this area ratio:

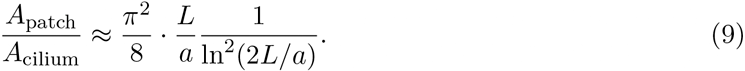

The area ratio as a function of the aspect ratio *L/a* is shown in Fig. 2b. For a typical cilium aspect ratio of *L/a* = 80 (with *L* = 10 *μ*m, and *a* = 125 nm), this area ratio is 3.8, implying that the cilium is much more effective per unit area than a receptor on the surface of the cell. Using an exact numerical result for the capacitance of a cylinder [32], the ratio becomes 4.5. With the dimensions given above, the radius of the circular patch with the same capture rate is *R* = 3.4 *μ*m.

### B. Cilium in shear flow

At the scale of cilia, the flow is characterised by a low Reynolds number, meaning that viscous forces dominate over inertia. The fluid motion is well-described by the Stokes equation, together with the incompressibility condition:

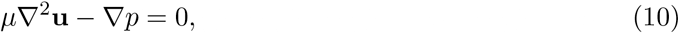

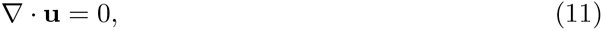

in which **u** is the fluid velocity, *μ* is the dynamic viscosity, and *p* is the pressure. The concentration field of some chemical species suspended within this fluid is governed by the advection-diffusion equation:

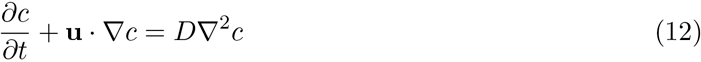

where *c* is a function of both position and time.

The ratio of advection to diffusion is described by the dimensionless Péclet number. This is usually written as some characteristic flow speed multiplied by some characteristic length scale, all divided by the diffusion constant.

Because the cilium grows from a flat surface with a no-slip boundary condition, the flow can be described as a uniform shear flow with the shear rate 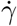. To estimate the capture rate constant of a cilium in a shear flow, we make use of the fact that the radius of the cylinder is much smaller than the length scale over which the shear flow varies. We therefore approximate the local rate density at any point on the cilium with that of an infinitely long cylinder in a uniform flow with velocity 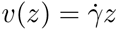. Due to Stokes’ paradox, there is no solution for the flow around a long cylinder at Re = 0, but the flow around a finite cylinder can be well described by assuming a small, but finite, Reynolds number. With Re = 0.2, the capture rate per unit length is [33]:

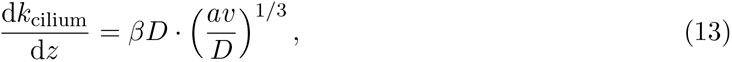

where *β* = (0.632 × 2*π*) is a numerical constant. The total rate constant is obtained by integration over the cilium length

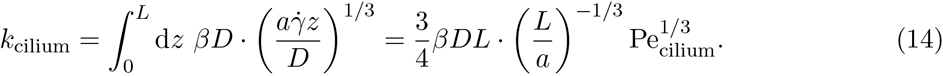

We take the characteristic velocity to be the speed of the cilium’s tip relative to the surrounding fluid, and hence the Péclet number for the extended cilium is

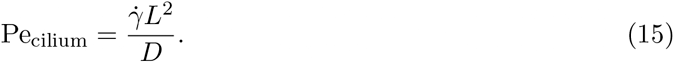

This expression for the rate once again shows a strong positive relationship between cilium length and sensitivity, as is known to be the case in real biological systems [31]. The characteristic Péclet number for the cross-over between the diffusive and the convective capture is of the order ~ *L/a* ≈ 80.

Once again we determine the size of a circular surface patch offering an equivalent effectiveness to the cilium in a flow with the same shear rate (Fig. 2c). The high-Pe rate constant for a patch in a shear flow is [34]:

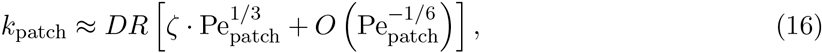

where 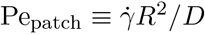 and *ζ* = 2.157 is a purely numerical constant. We can calculate the ratio of the area of the equivalent patch to the area of the cilium for these high-Pe asymptotic results:

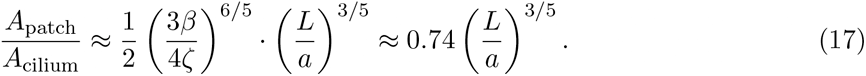

The area ratio is shown in Fig. 2d and and compared with the results in a quiescent fluid. For a typical cilium aspect ratio *L/a* = 80 (with *L* = 10 *μ*m, and *a* = 125 nm), this area ratio is 10 – much larger than the area ratio in a quiescent fluid, which was 3.8. The radius of the patch is 5 *μ*m. This means that a cilium is better per area than a patch at both low and high Péclet numbers, but the cilium excels when the Péclet number is large.

### C. Active pumping

A mounting collection of evidence suggests that both primary and motile cilia have sensory roles [20]. We are interested in the extent to which cilium motility can increase their ability to detect particles. To this end, we numerically simulate various different possible types of ciliary motion in otherwise quiescent fluids.

Because of the complex flow patterns and time-dependent boundary conditions, the absorption by a beating cilium is not analytically tractable. Instead, we use numerical simulations to determine the rate constants. We consider three different active pumping scenarios: a purely reciprocally moving cilium, a cilium tracing out a cone around an axis perpendicular to the surface, and a cilium tracing out a tilted cone (all shown in their respective order in Figs. 3a-c).

**FIG. 3:**
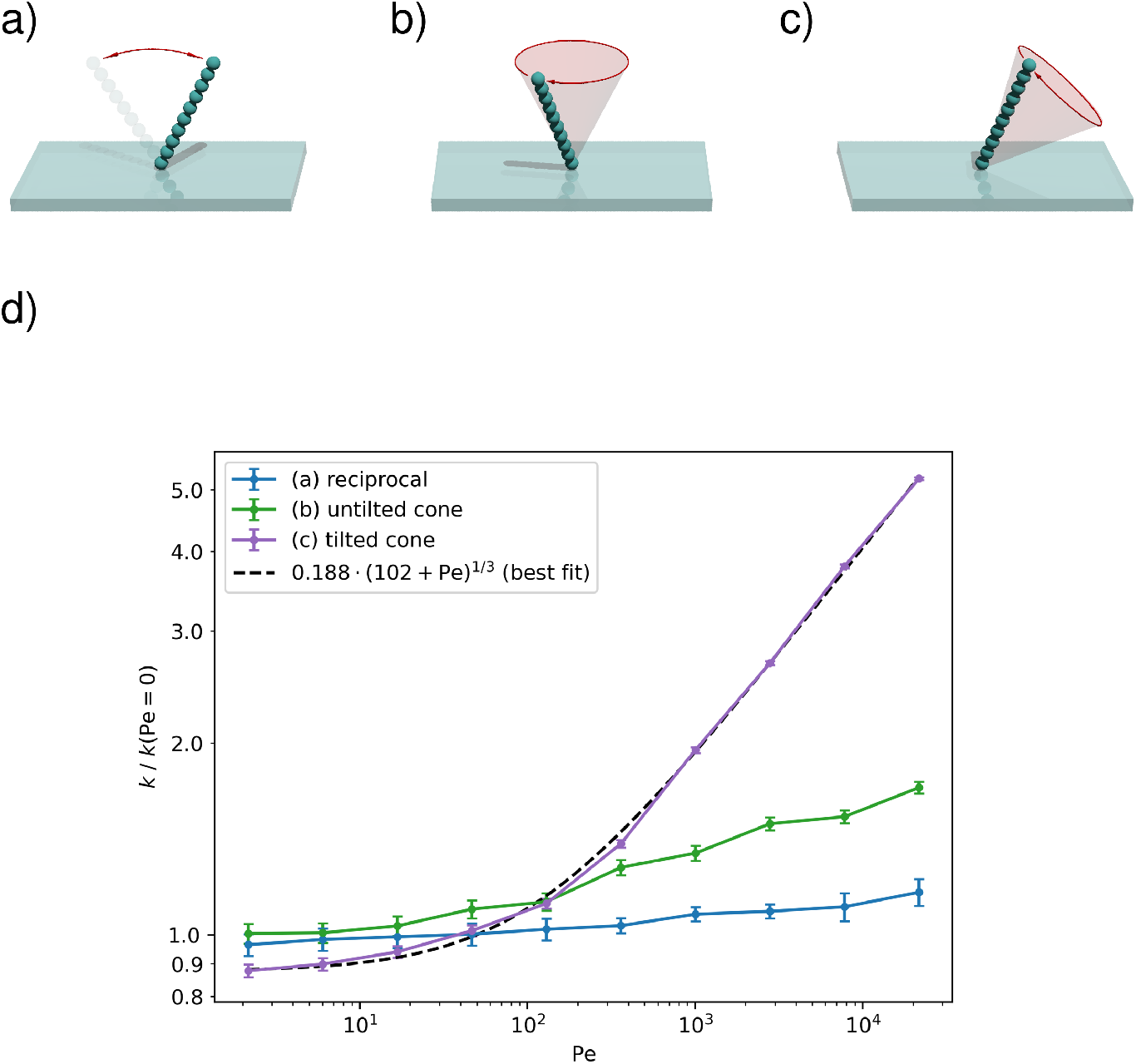
The capture rate on an active cilium for 3 types of motion. (a) The cilium is undergoing reciprocal motion, which is not generating any net flow. (b) The cilium moves along a cone with its axis perpendicular to the surface, such that it produces a rotational flow, but no long-range fluid transport. (c) The cilium moves along a tilted cone, which generates a long-range volume flow. (d) The capture rate constants *k* of a beating cilium as a function of the Péclet number. The rates are determined using stochastic simulations. The error bars denote 95% confidence intervals and the dashed line shows a fit function that interpolates between the high- and low-Péclet limits. All rates are normalised to the rate constant for a diffusion-limited capture with a cylindrical cilium with the same length and width.

The rate constants in these scenarios, relative to that of a non-moving cilium, are plotted in Fig. 3d. Analogously to the cilium in a shear flow, we define the Péclet number using the maximum tip velocity during the cycle:

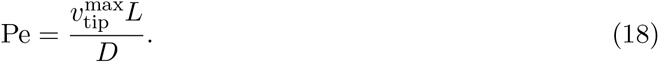

The reciprocally moving cilium (Fig. 3a) displays almost no improvement over several orders of magnitude of the Péclet number. This is expected, because Purcell’s scallop theorem states that purely reciprocal motion does not create any net flow, so the particle intake is largely diffusive in nature. A minor increase of the rate constant with the Péclet number is caused by the local shear flow that facilitates absorption on the surface.

The cilium moving around a vertical cone (Fig. 3b) induces a net rotational flow, but no inflow or outflow (by symmetry, the time-averaged flow can only have a rotational component [35]). Nevertheless, the constant motion of the cilium through the fluid leads to a higher local capture efficiency. The rate constant therefore shows more improvement; over a few orders of magnitude of the Péclet number, the rate constant increases by a factor of two.

The tilted cone (Fig. 3c) shows by far the highest capture rate, which is unsurprising. When the cilium is near to the plane, the no-slip boundary screens the flow, whereas when it is far from the plane, its pumping is unimpeded. This results in the cilium inducing a long range flow in one direction, characterised by a finite volume flow rate [36]. The long range flow causes a constant intake that replenishes the depleted particles. At high Péclet numbers, the capture rate scales *k* ~ Pe^1/3^, which is the same dependence as in an external shear-flow, although with a prefactor that is lower by a factor of ~ 2. Locally, the relative flow around the cilium is the same whether a cilium is pivoting or resting in a shear flow. The pumping effect of the tilted cilium, on the other hand, provides sufficient inflow that the concentration around a cilium sees only a small depletion effect.

### D. Collective active pumping

We consider seven cilia on a hexagonal centred lattice with lattice-constant 0.95*L*, with a view to understand how the presence of multiple cilia affects performance. We quantify the performance gain using a quantity *Q*, which we define as

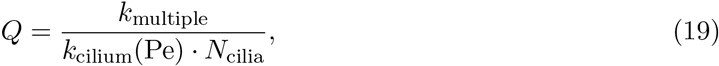

which represents the fractional per-cilium improvement in rate constant compared to a single isolated cilium at the same Péclet number.

Using numerical simulations, we find that at zero Péclet (Figs. 4a-b), *Q* ≈ 0.5, which means that the cilia locally deplete the concentration field, harming the per-cilium effectiveness; in a quiescent fluid, it is most efficient for cilia to stand far away from their neighbours.

**FIG. 4:**
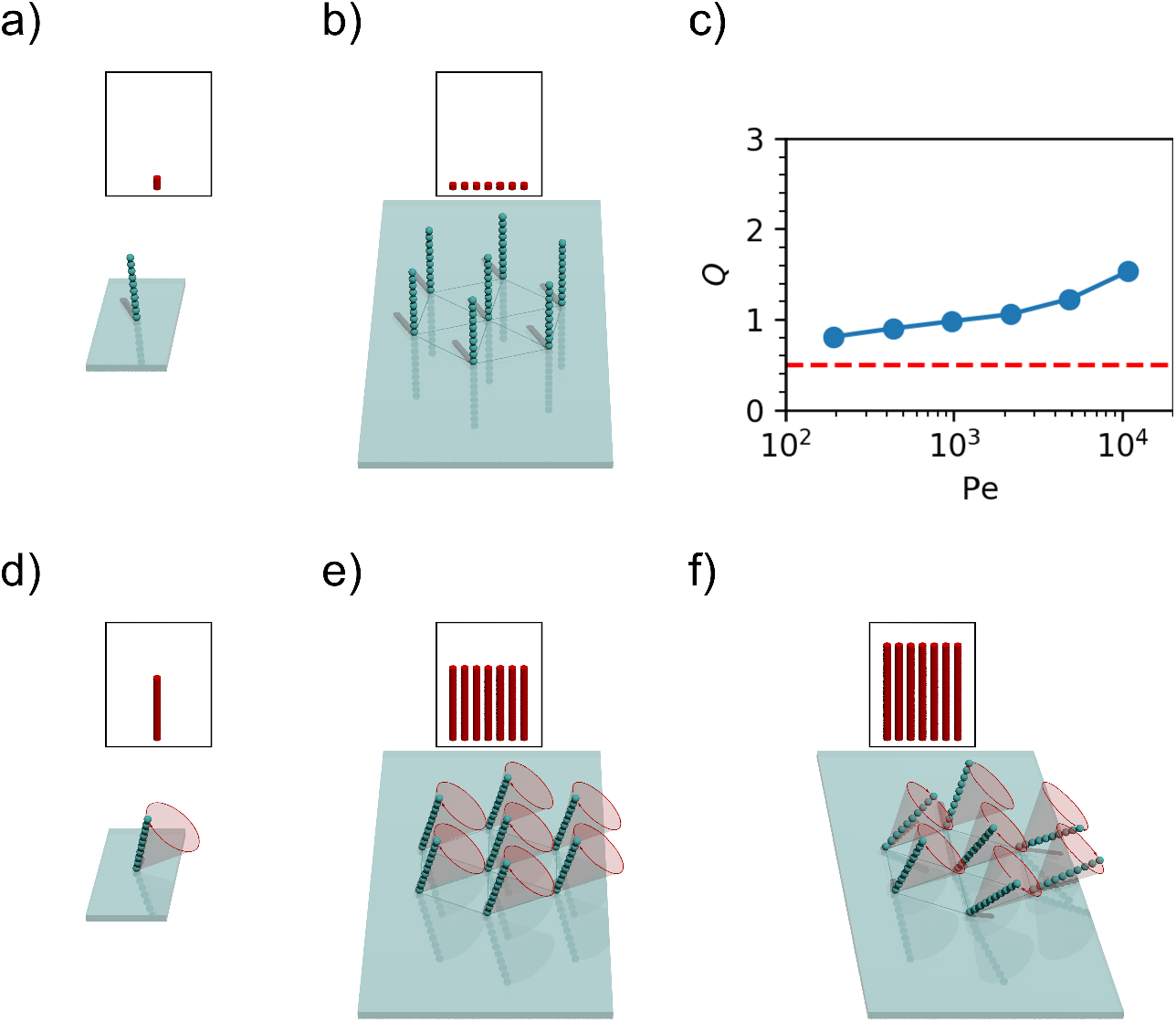
Comparison between the capture rate constant of a single cilium (a, d) and a bundle of *N*_cilia_ = 7 cilia (b, e, f). In the insets, the height of each red cylinder indicates the rate constant per cilium at Pe ≈ 10000, and the number of cylinders represents the number of cilia. For immotile cilia (a, b), a bundle has a lower per-cilium capture rate than an isolated cilium, although the the total rate constant of the bundle is higher. The reduced capture rate per cilium is caused by the depletion of ligands close to the bundle. For motile cilia (d, e, f), the situation is reversed and the capture rate per cilium in a bundle (e, f) can be significantly higher than for an isolated cilium (d). The increase can be explained by the collective flow generation, which helps the capture on all cilia. In (e) the cilia all beat with the same frequency corresponding to Pe ≈ 10000 but with identical phases. In (f) all cilia beat with the same frequency corresponding to Pe ≈ 10000, but their phases are chosen randomly. It can be seen that the random phases give a higher rate constant than the uniform phases. (c) shows how the performance gain *Q* varies with the Péclet number. The rates shown at each point are the average of 30 random phase configurations like the one shown in (f). The red line is the *Q*-value for Pe = 0.

However, when the cilia actively move (with each tracing out a tilted cone with a different randomly-chosen phase lag compared to its neighbours, as in Fig. 4f) the trend is reversed: we find that at Pe ≈ 10000, *Q* ≈ 1.53, meaning that per cilium, the capture rate is around 50% higher in the collective when compared to an isolated cilium with the same Péclet number. We find that over the range of Péclet numbers simulated, the *Q* increases monotonically with the Péclet number (see Fig. 4c).

When comparing these randomly chosen phases to a patch of cilia which beat in uniform, we find that cilia which beat in phase (Fig. 4e) see an improvement over the stationary case with *Q* ≈ 1.16, but are much less effective than the cilia patch that beats with random phases. The random phases give a higher volume flow, and complex hydrodynamic interactions between the randomly-phased cilia result in a slightly higher capture chance for any given particle.

## III. DISCUSSION

Our results address a simple question: why are so many chemical receptors located on cilia? Besides the well-known advantages of compartmentalisation, which facilitates the downstream signal processing, we show that the elongated shape of a cilium provides an advantage for the capture rate of molecules in the surrounding fluid. The advantages can be summarised as follows:

1. If neither the fluid nor the cilium move and the process of particle capture is purely diffusive, the elongated shape improves the capture rate of the cilium by giving it better access to the diffusing ligands. The length dependence of the capture rate has the sub-linear form *k* ~ *L/* log *L*. With typical parameters, the cilium achieves a capture rate equivalent to that of a circular patch of receptors on the epithelial surface with 4× the surface area of the cilium.
2. When a non-moving cilium is exposed to a shear flow, the advantage increases, mainly because the tip of the cilium is exposed to higher flow velocities. The capture rate scales with *k* ~ *L*^4/3^ and becomes equivalent to that of a surface patch with approximately 10× the surface area at high flow rates.
3. An actively beating cilium can achieve capture rates comparable to those by a passive cilium in a shear flow with the same relative tip velocity, but only if the beating is non-reciprocal, i.e., if the cilium generates a long range directed flow. The capture rate scales with the beating frequency to the power of 1/3.
4. Without motility, a bundle of sensory cilia achieves a capture rate *per cilium* that is lower that than of a single cilium, because of the locally depleted ligand concentration. However, the situation can become reversed if the cilia are beating: then each cilium benefits from the flow produced by the bundle as a whole, and the per-cilium capture rate can be significantly higher than in an isolated beating cilium. Cilia beating with random phases achieve significantly higher capture rates than when beating in synchrony.

Our results are based on a few assumptions. We assumed that the particles get absorbed and detected upon their first encounter of the cilia surface – an assumption that is justified if the receptors are covering the surface at a sufficient density [26], or if the particles bind non-specifically to the membrane of the cilium first. We also treat the particles as point-like (their size only has an influence on their diffusivity), which is accurate for molecules up to the sizes of a protein and we do not expect a significant error even for small vesicles. The Rotne-Prager tensor approximation used to determine the flow fields does not exactly satisfy the no-slip boundary condition on the surface of the cilium, especially at high Péclet.

With the typical dimensions of a cilium (*L* = 10 *μ*m, *a* = 0.125 *μ*m) and a diffusion constant of a small molecule *D* = 10^−9^ m^2^s^−1^, we obtain *k*_cilium_ = 0.7 pM^−1^s^−1^. A chemosensory cilium working at the physical limit is therefore capable of detecting picomolar ligand concentrations on a timescale of seconds. If the epithelium is embedded in mucous with a viscosity at least 3 orders of magnitude higher than water [37] (we disregard its viscoelastic nature here) and the molecule has a Stokes radius of a few nanometres, the diffusion-limited capture rate reduces to around *k*_cilium_ = 100 *μ*M^−1^s^−1^.

In a shear flow with a typical shear rate of 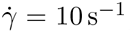, the Péclet number of a small molecule in water is of the order of Pe ≈ 1, where the capture rate still corresponds to the stationary case. However, with larger molecules and higher viscosities, the Péclet numbers can exceed 10^4^, leading to a significant enhancement of the capture rate.

When the same cilium is beating with a frequency of 25 Hz, the Péclet number is of the order ~ 10, which is too small to have an effect on the capture rate. Again, with larger molecules and higher viscosities, the Péclet numbers can exceed 10^5^, meaning that the motility accelerates the capture rate by one order of magnitude. Figure 5 shows how the molecular Stokes radius affects the fluid viscosity required to break the diffusion limit for a few different scenarios. However, when the particle size becomes comparable to the cilium diameter, the approximation that treats them as point particles loses validity. Indeed, it has been shown that particle size can have a direct steric effect on the capture rate [38]. Furthermore, the capture process of large particles can depend on a competition between hydrodynamic and adhesive forces [39]. Steric effects can even lead to particle enrichment in flow compartments [40].

**FIG. 5:**
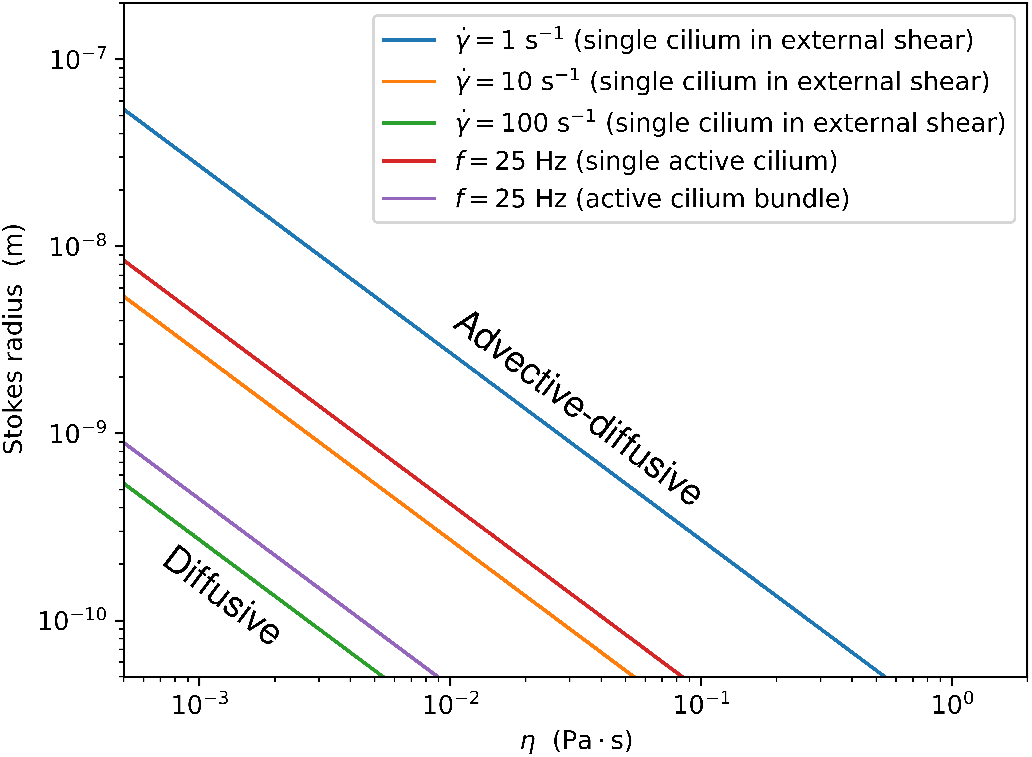
The demarcation between the regime where the rate constant is determined mostly by the diffusion limit and the regime in which it is enhanced by advection as a function of the fluid viscosity *η* and the particle Stokes radius. The blue, orange and green lines show the results for a passive cilium in a shear flow (Fig. 3c) and the magenta line for a bundle of 7 cilia (Fig. 4f). For all lines, the cilium dimensions are *L* = 10 *μ*m and *a* = 250 nm.

An example of a system in which active cilia can break the diffusion limit is the vertebrate left-right organiser (ventral node in mouse, Kupffer’s vesicle in zebrafish), where motile cilia drive a lateral flow that is sensed by other cilia. One hypothesis proposes that that the flow is sensed mechanically, by immotile crown cilia [41]. However, the mechanosensitive hypothesis has been challenged in Kupffer’s vesicle in zebrafish, which is one of the best characterised LROs. The paucity of immotile cilia at the mature stages of Kupffer’s vesicle, along with the weakness and high local variability of the flow, provide a strong argument against mechanosensation [6, 42], primarily because a mechanism that uses a motile cilium to sense a flow that is much slower than the cilium’s own motion appears unfeasible. The alternative proposition is that flow is detected via transport of signalling particles, possibly small vesicles termed “nodal vesicular parcels” [43]. According to this hypothesis, motile cilia also act as chemoreceptors. These cilia are approximately *L* = 6 *μ*m long and are exposed to typical shear rates in the fluid 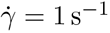 [42]. For a vesicle with a Stokes radius of 100 nm, the estimated Péclet number of a passive cilium in the shear flow is Pe = 17, too small to accelerate the capture rate. An active cilium beating at 25 Hz, on the other hand, has Pe = 1300. Although there are still many unknowns about the detection mechanism, we have shown that motile cilia can have capture rates that are several times higher than those of a passive cilium. Therefore chemosensitive detection seems to be a more plausible explanation of flow sensation in the left-right organiser, due to the benefits of combining motility and sensory function in the same cilium.

We have thus proven that the geometry of a cilium always means an advantage in chemical sensitivity over receptors on the epithelial surface, whether in a quiescent or moving fluid. At high Péclet numbers, which are achieved in viscous fluids, with very large particles or in very strong flows, the advantage of a cilium increases further. Motility also confers an increase in sensitivity, and this advantage is further pronounced in a bundle of cilia. These advantages can work in concert with others, such as avoiding charged surfaces and glycocalix on the cilium’s outside and the provision of a closed compartment on the inside. Further work might examine the extent to which motility benefits cilia in a fluid with bulk flow, or investigate the effect of metachronal waves on ciliary chemosensitivity. Finally, our results shed light on possible engineering applications for microfluidic sensing devices based on these ideas, e.g. using magnetic actuation [44–46].

## IV. METHODS

Numerically-simulated point particles are injected into a finite system containing a motile cilium, and move around due to advection (resulting from the motion of the cilium) and diffusion, until they either escape from the system or are absorbed by the cilium. The proportion of particles which are captured is used to compute a rate constant.

### A. Flow calculation

The hydrodynamics are computed using a modified Rotne-Prager mobility tensor **M** that accounts for the no-slip boundary. If there are *N* spheres of equal radius *R* in the simulation, each having a prescribed trajectory **r**_*i*_(*t*) and each acted upon by a force **F**_*i*_(*t*), then these forces must satisfy [44]

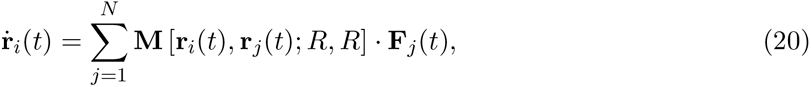

for every *i* ∈ [1, *N*]. Since every term except the forces is known, the forces can be determined numerically at a given *t* by solving this set of simultaneous equations. Then the fluid velocity at any point **x** can be determined by

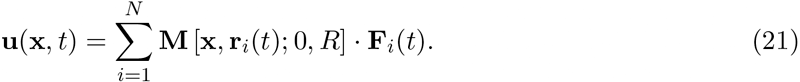

### B. Injection

We require a particle injection procedure that satisfies the concentration boundary condition *c* → *c*_0_ far from the absorbing cilium. We achieve this by introducing two bounding boxes in the simulation: an inner and an outer box, separated by a thin distance *d* (Fig. 6). The particles are injected at the boundary of the inner box and absorbed at the outer box. The injection rate is calculated such that it corresponds to the advective-diffusive flux through the layer between the boxes if the concentration at the inner box is *c*_0_. Because the flux through the boundary layer is much larger than the flux of particles absorbed inside the inner box, the method is suited to ensure a constant concentration boundary condition. The method is similar to a recent algorithm using a single boundary [47], but uses a simpler injection function.

**FIG. 6:**
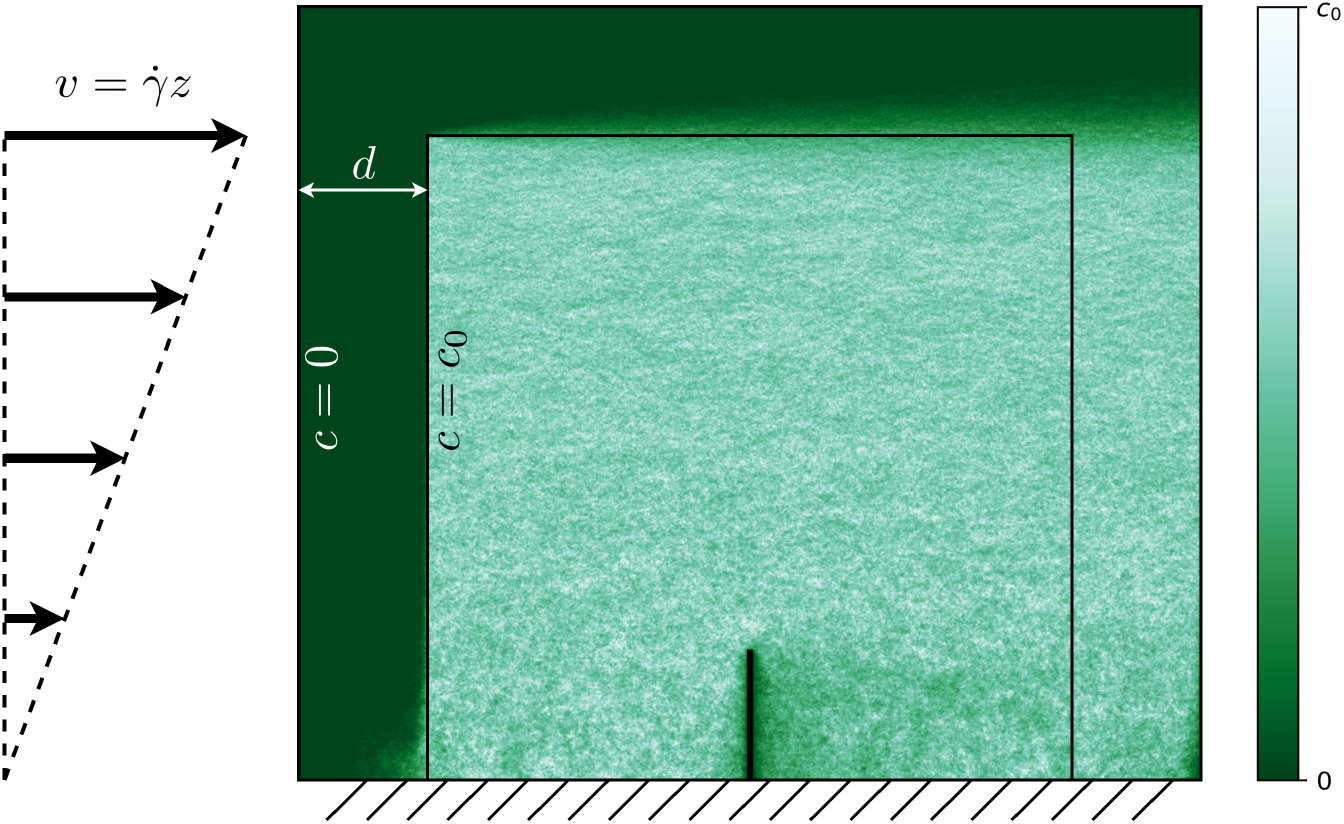
The boundary conditions used for injection. Since the fraction in incident particles absorbed by the cilium is small compared to the fraction absorbed by the outer surface, the concentration at the inner boundary is very close to *c*_0_. The coloured overlay shows the concentration as recorded in an example numerical simulation of a cilium in a shear flow with Pe = 50.

To calculate the injection current density, we solve the one-dimensional steady-state advection-diffusion equation

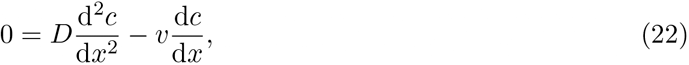

with the boundary conditions *c*(0) = 0 and *c*(*d*) = *c*_0_. The solution is

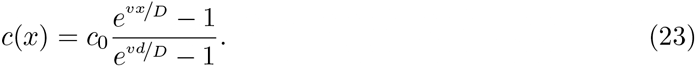

By the application of Fick’s law, this leads to an expression for the current density through the inner box:

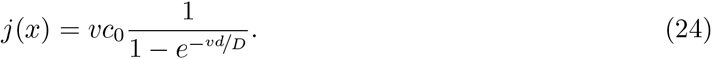

We assume that a test particle will take take much longer to reach the cilium than the characteristic time required for the flow to change, and hence we take 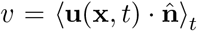, where 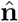 is the inward pointing surface normal of the inner box. This function can then be used to probabilistically weight where particles are most likely to be injected on the inner box.

### C. Numerical integration

The test particle position is updated using an Adams-Bashforth-Milstein multistep numerical integration method in the presence of noise [48]:

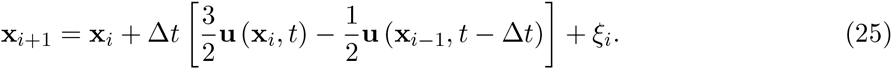

Because the computation of the flow field **u** (see Eq. 21) is the most demanding step, it is advantageous over methods that require additional function evaluations per step. *ξ_i_* is a vector where each element is pseudorandomly generated Gaussian noise with standard deviation 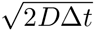 and mean of zero.

### D. Rate evaluation

We finish each simulation run when the particle position reaches the cilium (capture), or the outer box (escape). At the end, the rate constant is determined as

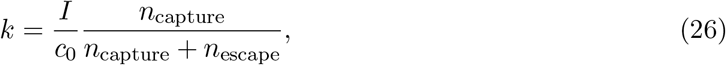

where *I* is the calculated total particle flux, obtained by integrating the flux density over the inner box, *I* = ∫ *j dS*.

### E. Numerical parameters

For all numerical simulations, we use a cilium consisting of 20 beads (thus giving a length to radius ratio *L/a* = 40). For the conical and reciprocal motion (Figs. 3a-c) we use an opening angle (between the cone axis and surface) of 30°, and for the titled conical motion (Fig. 3c) the axis of the cone is tilted relative to the vertical by an angle of 55°.

In the collective regime, the parameters are the same, with the addition of a hexagon lattice constant of 0.95*L*. The cones are tilted such that their axes are perpendicular to one chosen side of the hexagon (left to right in Fig. 4f).

## V. ACKNOWLEDGMENTS

We thank David Zwicker for comments on the manuscript. This work has been supported by the Max Planck Society. A.V. acknowledges support from the Slovenian Research Agency (grant no. P1-0099).

